# Novel mitochondrial genome rearrangements including duplications and extensive heteroplasmy could underlie temperature adaptations in Antarctic Notothenioid Fishes

**DOI:** 10.1101/2022.09.19.508608

**Authors:** Bushra Fazal Minhas, Emily A. Beck, C.-H. Christina Cheng, Julian Catchen

## Abstract

Mitochondrial genomes are known for their compact size and conserved gene order, however, recent studies employing long-read sequencing technologies have revealed the presence of atypical mitogenomes in some species. In this study, we assembled and annotated the mitogenomes of five Antarctic notothenioids, including four icefishes (*Champsocephalus gunnari, C. esox, Chaenocephalus aceratus*, and *Pseudochaenuchthys georgianus*) and the cold-specialized *Trematomus borchgrevinki*. Antarctic notothenioids are known to harbor some rearrangements in their mt genomes, however the extensive duplications in icefishes observed in our study have never been reported before. In the icefishes, we observed duplications of the protein coding gene *ND6*, two transfer RNAs, and the control region with different copy number variants present within the same individuals and with some *ND6* duplications appearing to follow the canonical Duplication-Degeneration-Complementation (DDC) model in *C. esox* and *C. gunnari*. In addition, using long-read sequencing and k-mer analysis, we were able to detect extensive heteroplasmy in *C. aceratus* and *C. esox*. We also observed a large inversion in the mitogenome of *T. borchgrevinki*, along with the presence of tandem repeats in its control region. This study is the first in using long-read sequencing to assemble and identify structural variants and heteroplasmy in notothenioid mitogenomes and signifies the importance of long-reads in resolving complex mitochondrial architectures. Identification of such wide-ranging structural variants in the mitogenomes of these fishes could provide insight into the genetic basis of the atypical icefish mitochondrial physiology and more generally may provide insights about their potential role in cold adaptation.

## Introduction

Mitochondria (mt) are specialized cytoplasmic organelles that provide substantial energy to eukaryotic cells and have enabled the evolution of eukaryotic complexity (Sagan, 1967). They contain their own genomes and are involved in significant biological processes like aerobic metabolism, stress response, energy balance, and oxidative phosphorylation (OXPHOS), among many others (Fontanesi, 2015; McBride et al., 2006). A typical metazoan mitogenome is very small (15-19 kilobasepairs (kbp)) having transferred many genes not needed for local metabolic control to the host organism’s nuclear genome (*i.e*. NuMTs, or nuclear mtDNA (Leister, 2005)). With few exceptions they possess a double-stranded, circular DNA molecule containing 13 protein coding genes, 22 transfer RNA (tRNA) genes (required for translation of proteins encoded by the mitochondrial genome (Boore, 1999)), two ribosomal RNA genes (*16S* and *12S*), a light strand origin of replication, and a control region (CR), which harbors transcription promoters and replication origins. Teleost mitogenomes in general harbor the same characteristics as typical Metazoan mitogenomes, but deviations have been identified involving duplications, local position changes (shuffling), and transpositions, while inversions are considered relatively rare (Gong et al., 2013; Inoue, 2003; Kong et al., 2009; Miya & Nishida, 1999; Satoh et al., 2016; Shi et al., 2013). Apart from gene duplications and insertion/deletion events, the majority of mt genome size variation is attributed to length differences in the CR, which might arise due to variability in the length or number of simple sequence repeats (Lee et al., 1995; Mignotte et al., 1990; Pereira, 2000).

Differing mitochondrial complements, called heteroplasmy, are detectable when mitogenomes possess differences in their length or nucleotide sequence and are known to arise because of somatic mutations, paternal leakage, or biparental transmission (reviewed in Breton & Stewart, 2015). Somatic mitogenomic mutations are particularly prevalent as the mutation rate of mt genomes is roughly 5-10 times that of the nuclear genome (Brierley et al., 1998; Haag-Liautard et al., 2008; Itsara et al., 2014). The level of heteroplasmy can vary at different organizational levels within the individual (cells, tissues, and organs), and at a population level where different rates of heteroplasmy may be present in different individuals (reviewed in Parakatselaki & Ladoukakis, 2021; reviewed in Stewart & Chinnery, 2015). Heteroplasmy is biologically important as it results in the presence of a dynamic pool of mt genomes in an organism; it can be sustained as a result of (1) random genetic drift causing an increase in the population of a particular type of mitochondrial genome through an unbiased transmission to daughter cells (Stewart & Chinnery, 2015), (2) relaxed replication where the proportion of any mt genome variant can also increase as mitochondria are replicated and destroyed continually in non-dividing cells (Stewart & Chinnery, 2015), or (3) positive selection of a variant having a functional advantage (reviewed in Klucnika & Ma, 2019).

Antarctic notothenioid fishes – cryonotothenioids – are the principal group of teleost fishes endemic to the Southern Ocean (SO) (Eastman, 2005). During the last 40 million years, as the region has undergone climatic changes resulting in extreme cold environments, cryonotothenioids emerged as the dominant marine teleost taxon, having evolved various physiological and morphological adaptations (Eastman, 1993) the most well-studied being the antifreeze glycoproteins, which prevent the growth of ice crystals within the fish (DeVries & Cheng, 2005). Icefishes (Channichthyidae), the most derived cryonotothenioids, are further specialized for the cold (Sidell & O’Brien, 2006), and notably are the only vertebrates that lack hemoglobin and the active oxygen transport it provides (Cocca et al., 1995; Ruud, 1954) which has been compensated for by major cardiovascular changes (Fitch et al., 1984; Hemmingsen et al., 1972; Kock, 2005). Unique mitochondria physiology has also been identified in this group of fishes. The heart and muscle cells in icefishes contain high densities of mitochondria, and the presence of abundant organellar lipid membranes could facilitate oxygen flux into the cell and the mitochondrial matrix, given O_2_ is much more soluble in lipids than in aqueous cytoplasm (O’Brien & Sidell, 2000). The icefish mitochondria have been described as having a “unique form and function” for their special architectural features and activities (O’Brien & Mueller, 2010) even when comparing to the related, red blooded cryonotothenioids.

Despite these radical specializations for life in the cold, one species of icefish, *Champsocephalus esox*, migrated within the last two million years to warmer Patagonian waters notwithstanding the lower oxygen concentrations (Kock, 2005; Stankovic et al., 2002). This species represents a model of *adaption following adaption*, exhibiting physiological changes for life in a temperate environment originating from an already derived icefish physiology. *C. esox* displays both higher mitochondrial densities and inner membrane morphology different from that of other icefishes (Johnston et al., 1998; O’Brien & Mueller, 2010). Positive selection has also been observed to act on several nuclear genes related to mitochondrial function and morphology when compared to its Antarctic sister species, *Champsocephalus gunnari* (Rivera-Colón et al., 2022). These patterns of selection, combined with the observed mitochondrial phenotypes, suggest that the organelle plays a fundamental role in the adaptation to warmer, temperate environments.

Changes to the mitochondria in cryonotothenioids are not limited to icefish. The cryopelagic red-blooded bald notothen, *Trematomus borchgrevinki*, is a more basal notothenioid that forages in the platelet ice layer under surface fast ice of McMurdo Sound, the coldest and iciest habitats in the SO. While stenothermal and adapted for extreme cold, the bald notothen is known to have retained some amount of plasticity in response to heat stress (Bilyk & DeVries, 2011). The mitogenome of *T. borchgrevinki* has been reported to harbor a large inversion (Papetti et al., 2021); however, the genetic details behind this inversion, and its role (or lack thereof) in cold specialization, remains unclear.

While many specific phenotypic changes for survival in the extreme cold have been described in cryonotothenioids, the molecular mechanisms of cold adaptation are not completely understood. Moreover, the role of the mitogenome in adaptive evolution remains little explored, however there are various studies which highlight the significance of mt genomic components in compensating for changing environments, including the role of *ND6* in high altitude adaptation in Tibetan horses (Xu et al., 2007), selection on the *ND4* and *ND5* genes to adapt to an active marine lifestyle in sea turtles (Ramos et al., 2020), and association of the *ND5* gene with cold tolerance in Chinese tiger frogs (Jin et al., 2022).

Most of the mt genomes available today were generated by long-range PCR coupled with first-generation Sanger sequencing, or directly using second-generation short-read sequencing. Both the short lengths of second-generation reads (100-300 bp) and limitations in PCR are unable to correctly resolve the repetitive sequences of the mt CR (Hommelsheim et al., 2015; Rayamajhi et al., 2022) or identify structural changes across the mitogenome. With the availability of third-generation, long-read sequencing the entire mt genome can be captured in a single read enabling assembly without any ambiguity. Recent studies using long-read sequencing have discovered spans of tandem repeats within the control region of mitogenomes (Kinkar et al., 2020, 2021). Any of these mt elements that are longer than the insert length of short-reads will be unresolvable by a short-read assembler, including inversions and tandem duplications (Rayamajhi et al., 2022).

In this study, we analyze the assembly and architecture of the mitogenomes of five cryonotothenioids: four icefishes (*Champsocephalus gunnari, Champsocephalus esox*, *Chaenocephalus aceratus*, and *Pseudochaenuchthys georgianus*) belonging to family Channichthyidae, and *Trematomus borchgrevinki* from the red-blooded subfamily Trematominae, using long-read sequencing (Fig. 1A). This is the first report on the mitogenome of the secondarily temperate icefish *C. esox* and the first complete long-read assemblies of the other icefishes and *T. borchgrevinki*. We found the mitogenomes of cryonotothenioids have undergone extensive rearrangements, with icefishes exhibiting tandem duplications of a region containing the *ND6* gene, *trnE* and *trnP* tRNAs, and the CR (hereafter, *ND6*/*trnE*/*trnP*/CR), and *T. borchgrevinki* displaying a large inversion, first reported by Papetti et al., (2021), that contains a set of CR tandem repeats. We also identified potential evidence of the duplication-degeneration-complementation (DDC) model (Force et al., 1999) in action in *ND6* in *C. esox* and *C. gunnari* revealing the potential of these fishes as new evolutionary mutant models (Albertson et al., 2009; Beck et al., 2022) for studying OXPHOS and other mitochondrial-driven processes.

**Figure 1.**
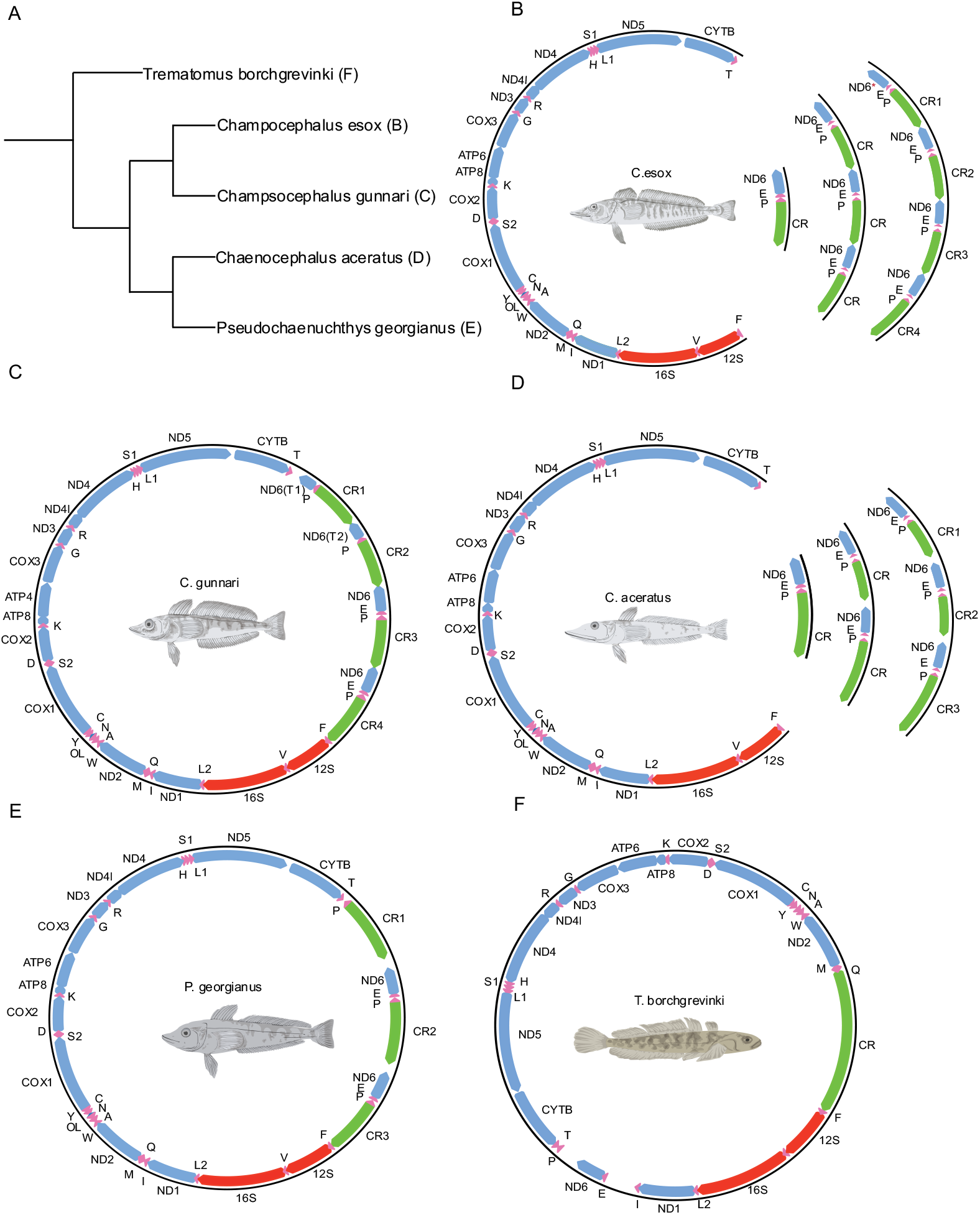
Complete annotated mitogenomes of species in this study. Protein coding genes are colored as blue, tRNAs as pink, ribosomal subunits genes as red, and control region is colored in green. (A) Phylogenetic relationship of fish in our study as shown in Near et al. (2018). (B) Mitogenome of *C. esox* showing heteroplasmy where we have observed reads showing variable mitogenomes with different numbers of tandem duplicated region *ND6*/*trnE*/*trnP*/CR, with reads having one, three and four copies of the duplicated block. (C) Mitogenome of *C. gunnari* with two complete blocks of *ND6*/*trnE*/*trnP*/CR region and two blocks with *ND6-T*/*trnP*/CR where *ND6-T* depicts truncated copy of *ND6* gene. (D) Mitogenome of *C. aceratus* showing heteroplasmy with one, two and three copies of the *ND6*/*trnE*/*trnP*/CR region. (E) Mitogenome of *P. georgianus* with two copies of duplicated *ND6*/*trnE*/*trnP*/CR region, and an additional copy of just *trnP*/CR. (F) Mitogenome of *T. borchgrevinki* showing an expanded control region within an inversion of 6551bp.

## Materials and Methods

### Mitochondrial sequence reads sources and sample preparation

The collection, handling, and tissue sampling of *C. gunnari, C. esox, and T. borchgrevinki* complied with University of Illinois, Urbana-Champaign IACUC approved Animal Use Protocols 07053 and 17148. We obtained mitochondrial sequences for the five selected cryonotothenioids from the whole genome raw read datasets of the respective species, summarized in Table 2 and detailed below.

**Table1.**
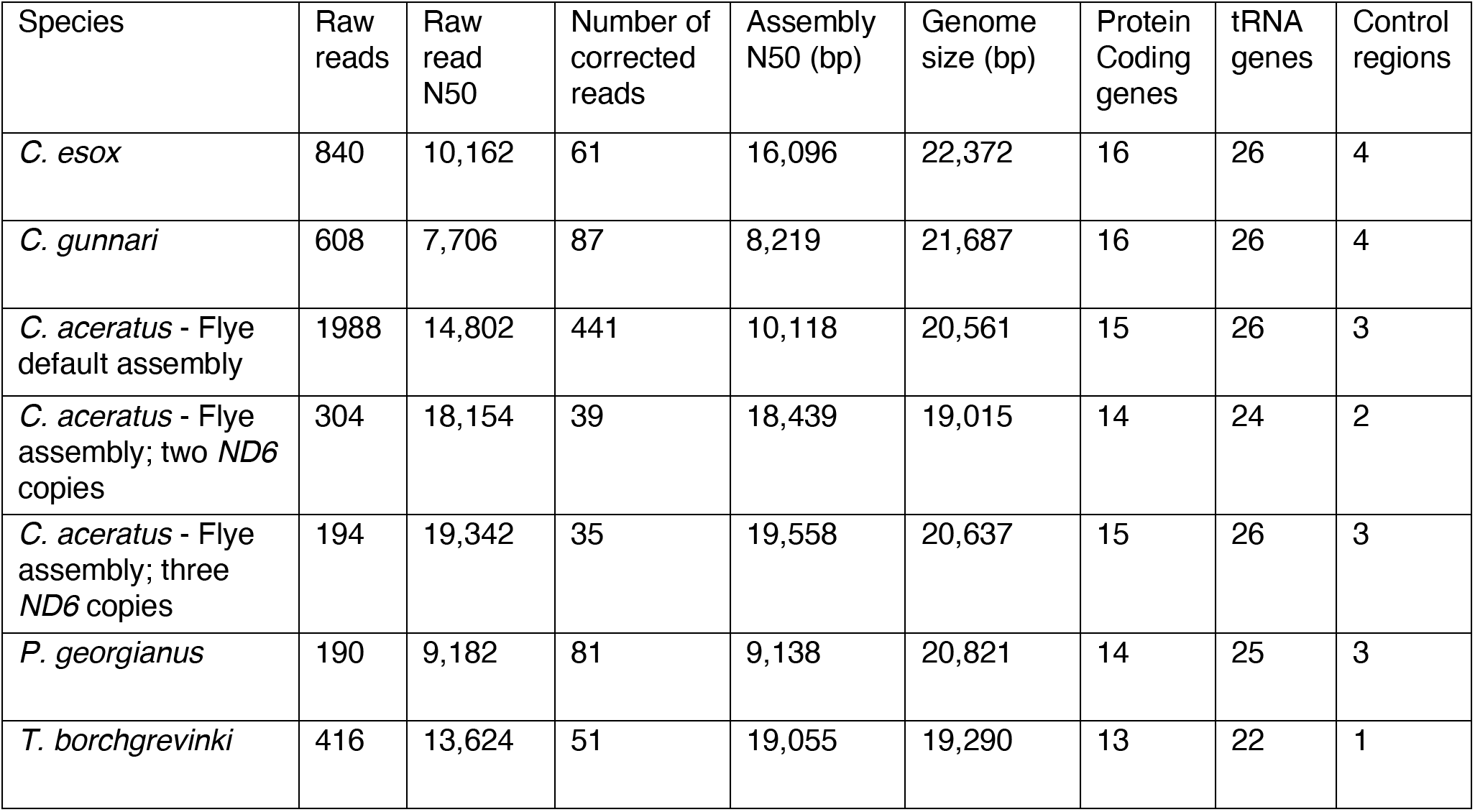
Raw read statistics, genome size and content of mt genomes of species in this study

**Table 2.** Sample information, sequencing technology and N50 raw read length for species sequenced in this study.

| Species | Sample location | Sample tissue | Technology | N50 |
| --- | --- | --- | --- | --- |
| *C. gunnari* (male)* | West Antarctic Peninsula | White muscle | PacBio Sequel II | 29.8 kbp |
| *C. esox* (male)* | Puerto Natales (Patagonia water) | Isolated hepatocytes | PacBio Sequel II | 24.3 kbp |
| *C. aceratus* (female) | King George Island, Antarctica | Muscle tissue | PacBio Sequel | 22.2 kbp |
| *P. georgianus* (female) | West Antarctic Peninsula | Spleen | PacBio Sequel | 9.8 kbp |
| *T. borchgrevinki* (female)* | McMurdo Sound, Antarctica | White muscle | PacBio Sequel II | 33.5 kbp |

For *C. gunnari* and *C. esox*, whole genome raw reads were generated by *de novo* sequencing for our companion genome projects. A single male specimen of *C. gunnari* caught from Gerlache Strait, West Antarctic Peninsula was sequenced using high molecular weight (HMW) DNA extracted from frozen white muscle. For *C. esox*, a single male specimen obtained from the Patagonia water near Puerto Natales, Chile was sequenced using HMW DNA derived from isolated hepatocytes. Methods of DNA preparation, Pacific Biosciences continuous long read (PacBio CLR) library construction, and sequencing on PacBio Sequel II instruments (2 SMRT cells each) are detailed in Rivera-Colón, et al., (2022). Briefly, sequencing yielded 10.7 million raw reads for *C. gunnari* with a mean and N50 read length of 29.7 kbp and 29.8 respectively. For *C. esox*, sequencing yielded 12.1 million raw reads with a mean and N50 read length of 13.1 kbp and 24.3 kbp N50 respectively.

For *C. aceratus*, we obtained the sequenced PacBio Sequel data available as BioProject PRJNA420419, from NCBI accessions SRR6942631 and SRR6942632. The data were derived from sequencing genomic DNA isolated from muscle tissue of a single female fish collected from Marian Cove, King George Island, Antarctica. The prepared genomic libraries were sequenced on PacBio Sequel System using P6-C4 sequencing chemistry (Kim et al., 2019). The BioProject provided 6.5 million raw reads, with a 13.6 kbp mean and 22.2 kbp N50 read length.

For *P. georgianus*, whole genome PacBio raw reads were available from NCBI under BioProject PRJEB19273, and accessions ERR3197127 and ERR3197122. The data were derived from sequencing DNA isolated from frozen spleen of a single female collected from the coast of Low Island, West Antarctic Peninsula. A PacBio CLR library was prepared using PacBio SMRTbell Template Prep Kit 1.0, and sequenced on a PacBio Sequel (Bista et al., 2022). The BioProject provided 7.4 million raw reads, with a 7.1 kbp mean and 9.8 kbp N50 read length.

For *T. borchgrevinki, de novo* sequencing was carried out for this study using a single female specimen caught from McMurdo Sound, Antarctica (78°S). HMW DNA was extracted from liquid nitrogen frozen white muscle using Nanobind Tissue Big DNA kit (Circulomics), lightly sheared for a ~75 kbp target, and used for PacBio CLR library construction. The library was selected for inserts ≥ 25 kbp using the Blue Pippin (Sage Science) and sequenced on one SMRT cell on PacBio Sequel II system using Sequel chemistry v.2 with 30 hours of data capture. The sequencing yielded 7.7 million raw reads, with 23.7 kbp mean and a 33.5 kbp N50 read length. Library construction and sequencing were carried out at the University of Oregon Genomics & Cell Characterization Core Facility (GC3F).

### Genome assembly and annotation

For all five notothenioids, raw reads were mapped against available mt reference genomes using minimap2 (Li, 2018). *C. gunnari* and *C. esox* were both mapped against *C. gunnari* (NCBI accession NC_018340). The *C. aceratus* and *P. georgianus* raw reads were mapped to NCBI accessions NC_015654.1 and NC_057673.1, respectively, *T. borchgrevinki* was mapped against NCBI accession KU951144.1. For *C. esox, C. aceratus, and P. georgianus*, reads with a mt matching block of at least 5000 bp were extracted using seqtk (https://github.com/lh3/seqtk) in order to assemble the mitogenome. For *C. gunnari* and *T. borchgrevinki*, reads with a matching block of at least 3000 bp were extracted the same way. Each set of raw reads (Table 1) were corrected using the corrections module of the Canu 1.8 assembler (Koren et al., 2017), and the corrected reads were then assembled *de novo* using the Flye assembler (v2.7) (Kolmogorov et al., 2019). The genomes were then annotated using Mitos2 (Donath et al., 2019), and tRNAscan-SE 2.0 (Chan et al., 2021). To avoid any inconsistencies and ambiguities, the annotations were manually checked using NCBI blastn (Camacho et al., 2009). We were unable to annotate the origin of replication for light strand (OriL) for *T. borchgrevinki*. Thus we searched for it around its canonical location, and by using RNAstructure (Reuter & Mathews, 2010) we identified a region with a hairpin structure, which we assigned as the putative OriL. The non-coding regions were explored for putative repeats using Tandem Repeat Finder (TRF) (Benson, 1999).

### K-mer analysis

The icefish mt genome assemblies indicated tandem *ND6* gene duplications. To confirm the assembly results and analyze the genic architecture of this region of the mitogenome we used k-mer analysis, which would allow us to search for *ND6* genes directly within the raw reads while allowing for sequencing errors. For each icefish we used the annotated *ND6* gene as query and the raw reads as subjects of the search; we k-merized both and searched for blocks of matching k-mers. To find the number of *ND6* copies and avoid random k-mer matches, we used the number of nucleotides between consecutive blocks of k-mer matches as a threshold for defining the start and end of putative *ND6* genes. For the k-mer size 19, we set the threshold of 800 nucleotides, that is if two matching k-mers are more than 800 nucleotides apart, they are considered parts of different *ND6* genes.

The variable length of raw, PacBio long-reads was problematic in visualizing the number of copies of *ND6* genes per read. To avoid the problem of variable read lengths, which may encompass different subsets of mt genes, we extracted reads containing both the *12S* and *CYTB* gene boundaries enclosing the *ND6*/*trnE*/*trnP*/CR tandem duplicated block, and then calculated the number of gene copies present in reads that spanned this full region. In the case of *C. aceratus* and *C. esox* our k-mer analysis indicated the presence of different numbers of *ND6* genes in different reads. For *C. aceratus*, apart from the generation of a primary assembly as discussed above, we separated the raw reads containing two or three putative *ND6* copies (Table 1) and assembled mt genomes for them independently using the same methods as explained above. For *C. esox*, however we did not have enough reads for each mt genome variant needed by the assembler to assemble the complete mitogenomes separately.

### Phylogenetic Analysis

The *ND6* sequences and control regions from all the icefishes were aligned with those of the basal, non-Antarctic notothenioid *Eliginops maclovinus* as an outgroup (Cheng, et al. in prep). We used Geneious 2022.1.1 (https://www.geneious.com) and aligned the sequences using MUSCLE (Edgar, 2004) with default parameters. The alignment was used to make a phylogenetic tree using PhyML (Guindon et al., 2010) using default settings with automatic model selection (BIC) and the tree was visualized in Figtree (http://tree.bio.ed.ac.uk/software/figtree/).

### Protein Structure Predictions

Protein sequences of ND6 from human (*Homo sapien*) and zebrafish (*Danio rerio*) were obtained from Ensembl version 107 (Cunningham et al., 2022) and compared to ND6 sequences from *C. aceratus, P. georgianus, C. esox*, and *C. gunnari* in this study, which contain *ND6* gene duplications. Full length ND6 amino acid sequences for each species and truncated ND6 sequences from *C. esox* and *C. gunnari* were run through the DeepTMHMM (TransMembrane Hidden Markov Model) to predict protein structure (Hallgren et al., 2022).

## Results

### Mitogenome assembly of *Champsocephalus esox*

The mt genome for *C. esox* has been reported for the first time in this study and is 22,372 bp in length, containing 16 protein coding genes, 28 tRNA genes, two ribosomal subunit genes, and four CR sequences (Table 1). The increased number of genes is due to the presence of a duplicated *ND6*/*trnE*/*trnP*/CR segment in tandem four times. Out of the four control regions, the first three are about the same length as typical control regions (1004, 1001, and 1004 bp), whereas the last control region is 1,097 bp in length, and the CRs did not contain any repeats. This duplication also results in four copies of the *ND6* gene along with intact *trnE* and *trnP* tRNA genes (Fig. 1B). Out of the four copies of *ND6*, one has a single base insertion, which shifts the reading frame and results in a truncated translated protein 81 amino acids in length that is different from the other three complete ND6 proteins, each 175 amino acids in length. The wildtype full length ND6 protein mirrors the structure of that seen in humans and other traditional animal models like zebrafish (Fig. 2A; Fig. 2B; Fig. 2E; Fig. S1). Recent predictions using deep learning models predict a 5 transmembrane domain (TMA-E) protein which differs from the original prediction of 6 TMs (Chinnery et al., 2001)(Fig. 2; Fig. S1). The truncated copy in *C. esox* terminates near the midpoint of the full wildtype protein sequence, retaining wildtype structure of the N-terminal half of ND6 through TMC (Fig. 2F, S1).

**Figure 2.**
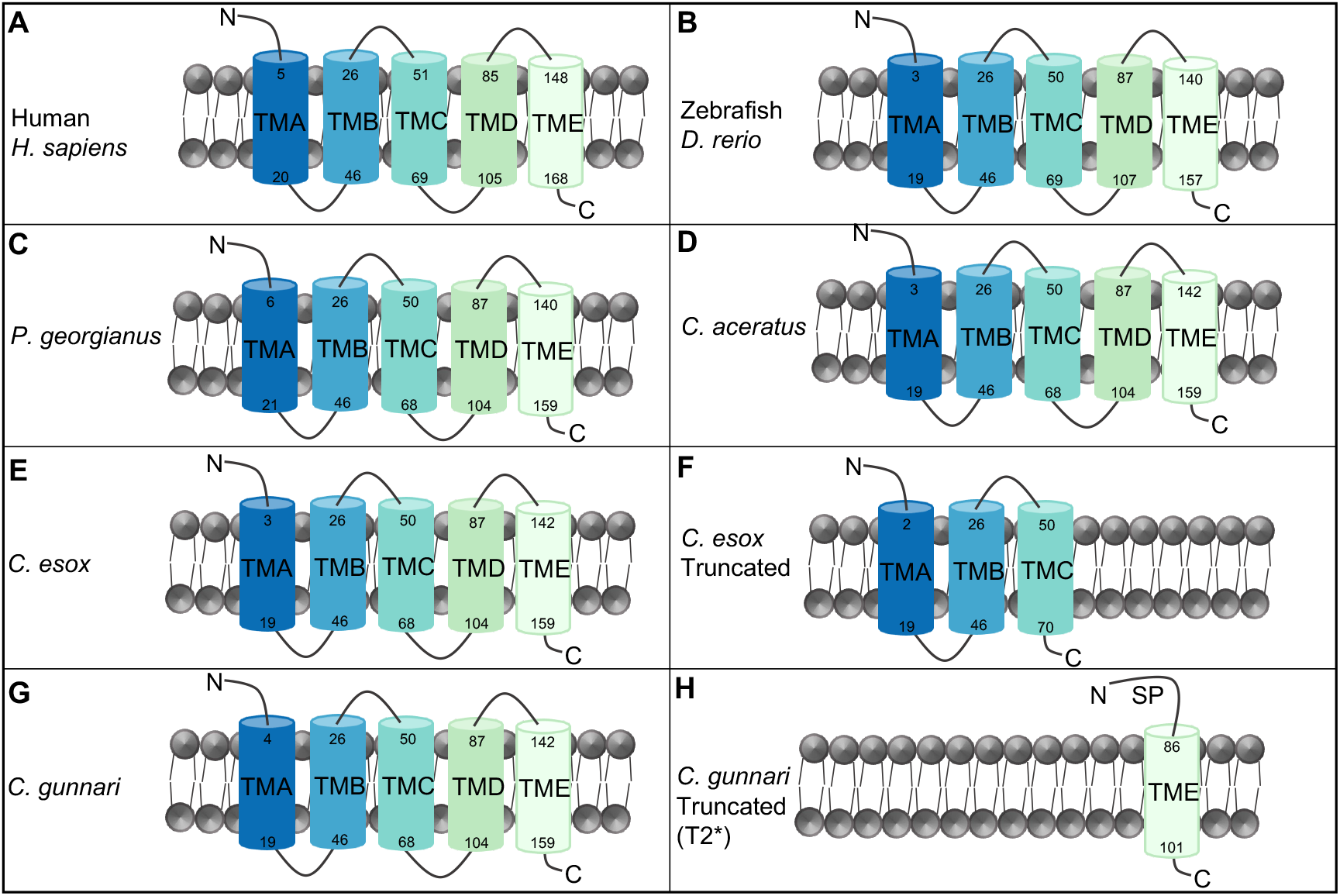
Protein structure of complete ND6 proteins contain five transmembrane proteins (TMA-TME). Protein structure of a complete ND6 protein in (A) Human, (B) Zebrafish, (C) *P. georgianus*, (D) *C. aceratus*, and (E) wildtype *C. esox*. (F) Protein structure of a truncated ND6 protein in *C. esox*, where the protein structure terminates near the midpoint of complete ND6 protein retaining wildtype structure of the N-terminal half of ND6 through TMC. (G) Protein structure of a complete wild type ND6 in *C. gunnari*. (F) Protein structure of truncated copy of ND6-T2 in *C. gunnari* where the C-terminal half with one transmembrane TME is retained along with predicted signal peptide (SP) at the N-terminal end. Only T2 truncation is shown as T1 is likely pseudogenized as the transcript is too short to allow for structure prediction.

Delving beyond the standard assembly, the k-mer analysis of raw reads enabled us to discover the presence of heteroplasmy in *C. esox* – i.e., a variable set of mitogenomes in this individual. There were 59 reads spanning the *12S* - *CYTB* block (the region containing the repeated *ND6*/*trnE*/*trnP*/CR block). Two of these reads had one *ND6* copy, 11 reads contained three *ND6* copies, 33 reads had four copies, one read had five copies, and the rest of the reads spanned the other side of the mitogenome. Along with *ND6* copies they also maintained the rest of the *ND6*/*trnE*/*trnP*/CR region. While various sequence (Macey et al., 2021) or length (Irwin et al., 2009) heteroplasmy in mt genomes have been reported, as far as we know, *C. esox* mt heteroplasmy is the most extensive case, with co-occurrence of different variants of mt genomes, each carrying variable numbers of genes (in variable numbers of duplicated blocks).

### Mitogenome assembly of *Champsocephalus gunnari*

The mt genome of *C. gunnari* is 21,687 bp in length, 2.8 kbp longer than the 18,863 bp reference. The genome consists of 16 protein coding genes, 26 tRNA genes, two ribosomal subunit genes, and four CRs (Table 1). The greater number of protein coding genes, tRNA genes, and CRs are due to a duplicated region containing *ND6*/*trnE*/*trnP*/CR four times in tandem (Fig. 1C), similar to *C. esox* but with important differences. Three of the four CRs have lengths of 1002, 1003, and 1004 bp, whereas the last CR is a larger sequence of 1,103 bp. Like *C. esox*, the CRs also did not contain any tandem repeats. In *C. gunnari*, however, we observed an alteration of the start site for *COX1* from an ATG to a GTG start codon and, more importantly, observed two complete *ND6*/*trnE*/*trnP*/CR duplicated blocks along with two additional blocks containing truncated copies of the *ND6* gene and a loss of *trnE* (Fig. 1C).

In both truncated copies of *C. gunnari ND6* there is an insertion in the same region as *C. esox* leading to a frameshift mutation, but *C. gunnari* also contains additional mutations upstream of the insertion resulting in alternative start sites, one in each truncated copy. In one truncated copy (T1), there is an insertion of four Gs and a transversion converting ATC to ATG. In this copy, the insertion of four nucleotides results in a frameshift leading to a possible transcript 69 bp in length. The second truncated copy (T2), however, contains an insertion of three Gs plus the ATC to ATG transversion retaining the open reading frame and a partially conserved C-terminal protein sequence of ND6 including TME (Fig. 2H).

Despite a highly derived mitogenome architecture, we did not find any heteroplasmy in the *C. gunnari* mitogenome.

### Mitogenome assembly of *Chaenocephalus aceratus*

The length of the primary *C. aceratus* mt genome assembly is 20,561 bp, 3.2 kbp longer than the reference (17,311 bp). It consists of 15 protein coding genes, 26 tRNA genes, two ribosomal subunit genes, and three CRs (Table 1). We observed three copies of the *ND6*/*trnE*/*trnP*/CR region present in tandem, in contrast to a single copy found in the reference genome. It appears the region underwent tandem duplication starting from the intergenic space between *trnT* and *ND6* and ending at the CR (Fig. 3A). We also observed a tandem duplication in one of the CRs, which is longer (1,306 bp) compared to the other two CRs which are both 847 bp (Fig. 1D).

**Figure 3.**
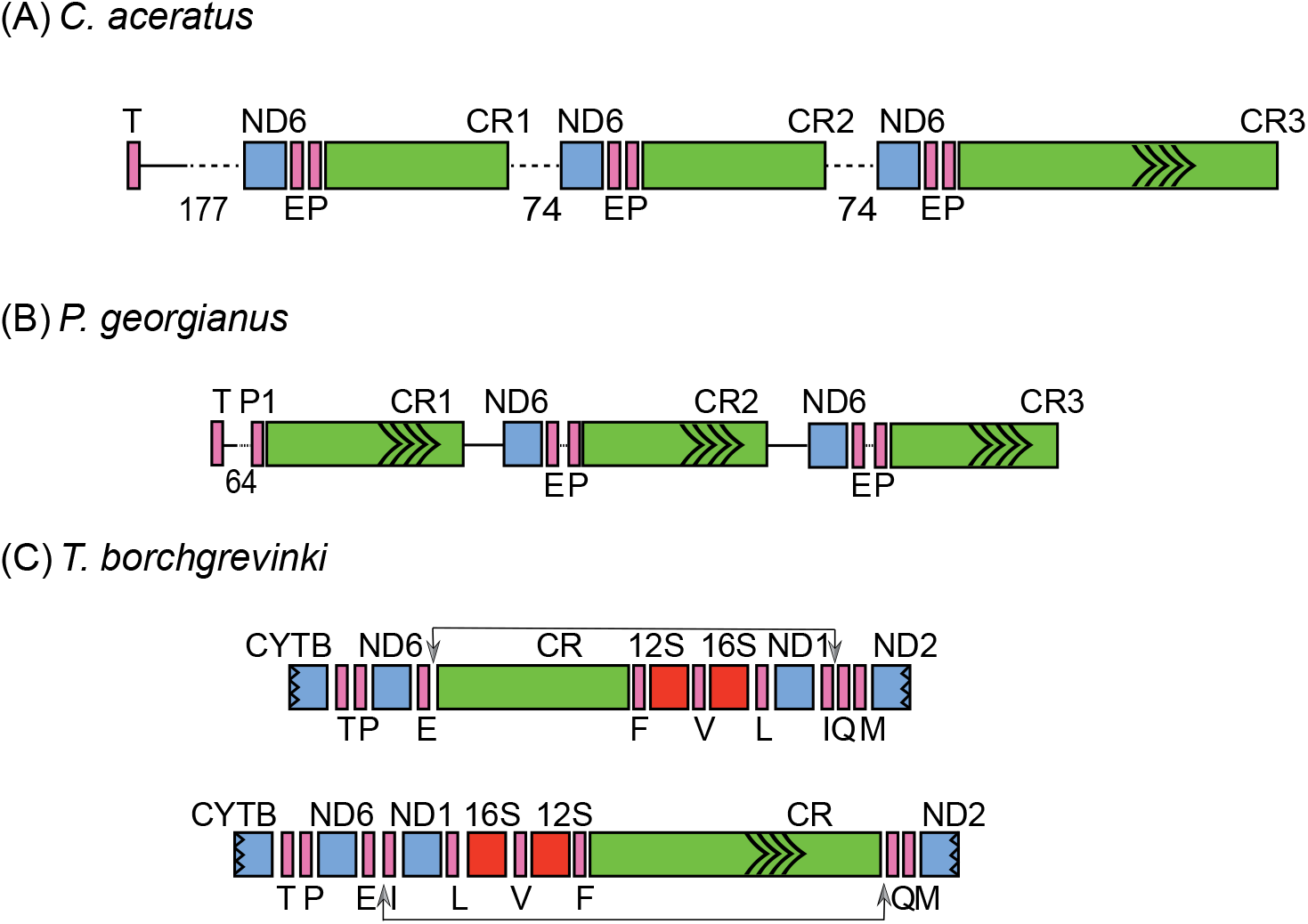
(A) Region showing the tandem duplicated block for C. aceratus with repeats in CR3. (B) Region containing the tandem duplicated block for P. georgianus with dotted region between T and P showing the potential remnant of the region between E and P, and all the control regions with repeats. (C) Comparison of the reference (top) vs our mt genome of T. borchgrevinki (bottom). The arrows on top show the position of the region which is inverted in our assembled genome. The arrows in the CR are the tandem repeats.

We observed heteroplasmy in *C. aceratus* using k-mer analysis, where we observed six reads having one *ND6* copy, 304 reads with two *ND6* copies, 193 reads with three *ND6* copies, and one read with four *ND6* copies. Each *ND6* copy occurs in a *ND6*/*trnE*/*trnP*/CR block, thus the variants with multiple *ND6* copies contain duplicated *ND6*/*trnE*/*trnP*/CR blocks. Two of the four element layouts were captured in enough reads to perform standard assemblies. These two sets of reads produced two different mt genomes, one mt genome assembly with length 19,015 bp and contained two copies of the *ND6*/*trnE*/*trnP*/CR block, while the other mt genome was 20,637 bp in length and recapitulated three copies of the *ND6*/*trnE*/*trnP*/CR region (Fig. 1D; Table 1).

### Mitogenome assembly of *Pseudochaenichthys georgianus*

The mt genome of *P. georgianus* is 20,821 bp in length, 3.5 kbp larger than the reference genome, which is 17,310 bp. The mitogenome contains 14 protein coding genes, 25 tRNA genes, two ribosomal subunit genes, and three CRs (Fig. 1E; Table 1). We observed two copies of the *ND6*/*trnE*/*trnP*/CR region present in tandem. We also observed an additional *trnP* and CR which seems to be the remnant of a third copy of the *ND6*/*trnE*/*trnP*/CR block. In other icefishes, *trnT* is followed by *ND6*, while in *P. georgianus*, it is followed by *trnP* instead. There is also an intergenic space between *trnT* and *ND6* that is of variable length (304 bp in *C. esox*, 197 bp in *C. gunnari*, and 177 bp in *C. aceratus*). In *P. georgianus*, the space between *trnT* and *trnP* is only 64 bp. When analyzed, a part of this region (from 38-64 bp) was identical to the intergenic region between *trnE* and *trnP*. The reduced length and the presence of the intergenic region between *trnE* and *trnP* might indicate the prior presence of a third copy of *ND6*/*trnE*/*trnP*/CR block, which may have been lost partially, removing most of the intergenic space between *trnT* and *ND6*, the *ND6* gene, and *trnE* while retaining the intergenic space between *trnE, trnP*, and the CR (Fig. 3B). The three CRs (two from the intact blocks and a third from one of the two remnant blocks) are larger in size compared to the typical CRs of fish mt genomes and are of variable lengths: 1439, 1377, and 1157 bp for CR1, CR2, and CR3 respectively. This variability in length is attributed to the presence of tandem repeats, where CR1 has a tandem repeat of 53 bp repeated 8.8 times, CR2 has repeat of 53 bp that is repeated 7.8 times, and CR3 has a repeat of 63 bp that is repeated 2.1 times. This is in stark contrast to *C. esox* and *C. gunnari* which do not contain tandem repeats.

Again, despite a highly derived mitogenome architecture, we did not observe any heteroplasmy in *P. georgianus*.

### Mitogenome assembly of *Trematomus borchgrevinki*

The architecture of the *T. borchgrevinki* mitogenome was distinct from the icefishes as well as the typical, teleost architecture. We assembled the longest mitogenome to date for *T. borchgrevinki* and within it we observed an inversion 6,551 bp in length along with tandem repeats within the control region. The mt genome of *T. borchgrevinki*, first reported by Liu et al., 2016 (NCBI accession KU951144.1; 17,299 bp), was assembled using Sanger sequencing and did not report any rearrangements. Later, Papetti et al., 2021 (NCBI accession MT232659.1; 18,325 bp) generated an additional mt genome of *T. borchgrevinki* using long-range PCR coupled with short-read sequencing and reported an inversion of at least 5,300 bp. Recently Patel et al., 2022 also generated a *T. borchgrevinki* mitogenome (NCBI accession MZ779011; 18,981 bp) using Illumina TruSeq synthetic reads (short-reads combined with a scaffolding technique) and found the same inversion, along with the presence of some intergenic spacer sequences. Using our PacBio long-reads we assembled a mitogenome of length 19,290 bp, which is 309 bp longer than the mt genome assembled by Patel et al., 2022, and the gene order and content are consistent with their findings. None of the previous assemblies reported the presence of tandem repeats in the mt genome of *T. borchgrevinki*.

Like the canonical vertebrate mt genomes, the *T. borchgrevinki* mitogenome contains 13 protein coding genes, 22 tRNA genes, along with two ribosomal subunit genes, and one CR (Fig. 1F; Table 1). Apart from *COX1* (which starts with a GTG codon), all protein coding genes use the ATG starting codon. The *CYTB, ND4*, and *COX2* genes do not have complete stop codons, which is a common observation in vertebrate mitogenomes, as they are known to be created via post-transcriptional polyadenylation (Ojala et al., 1981). The CR for this genome was substantially longer compared to its reference (i.e., 2,651 bp versus 1,212 bp) and is also substantially longer than the usual CR in teleost fishes. For example, the CR for zebrafish is only 950 bp (Broughton et al., 2001). The size of the CR was verified by examining the region between *12S* rRNA and *nd2* (which spans the CR) in raw PacBio reads. We found the length was consistent with our assembled CR and we did not observe any length heteroplasmy. The CR expansion is due to the presence of a high number of repeats which spanned 1,400 bp of the total length of the control region (2,651 bp). There were two sets of repeats, one was 291 bp long (spanning 157-448 bp in the CR) and contained three copies of a secondary tandem repeat of 97 bp. The second repeat block spans a region from 1,263 to 2,381 bp in the CR and contains additional nested repeats of variable length (Fig. 3C).

### Large Inversion in *T. borchgrevinki*

Similar to what was reported by Papetti et al., 2021, we also observed a large inversion in the mitogenome, though we found the inversion to be 1.2 kbp longer (6,551 bp in our data compared to 5,300 bp) (Fig. 1F). The inversion contains the CR that harbors the machinery for mitogenome replication. We manually assigned the putative OriL at the expected location (between the *trnN* and *trnC* tRNA genes; see Methods); it was considerably shorter in length (25bp) and did not form its usual hairpin structure.

### Evolution of duplicated genes in icefishes

To understand the evolutionary origin of the duplicated *ND6* copies and CRs, we performed a phylogenetic analysis of these genes and CRs with *E. maclovinus* as an outgroup. We observed that the *ND6* copies are more closely related within a species compared to their respective orthologs. Similarly, for CRs, the paralogs group together for each species showing that paralogs are more closely related than orthologs across species, except for the last CR of *C. gunnari* and *C. esox* which are longer than the first three CRs in both species and thus group together (Fig. 4).

**Figure 4.**
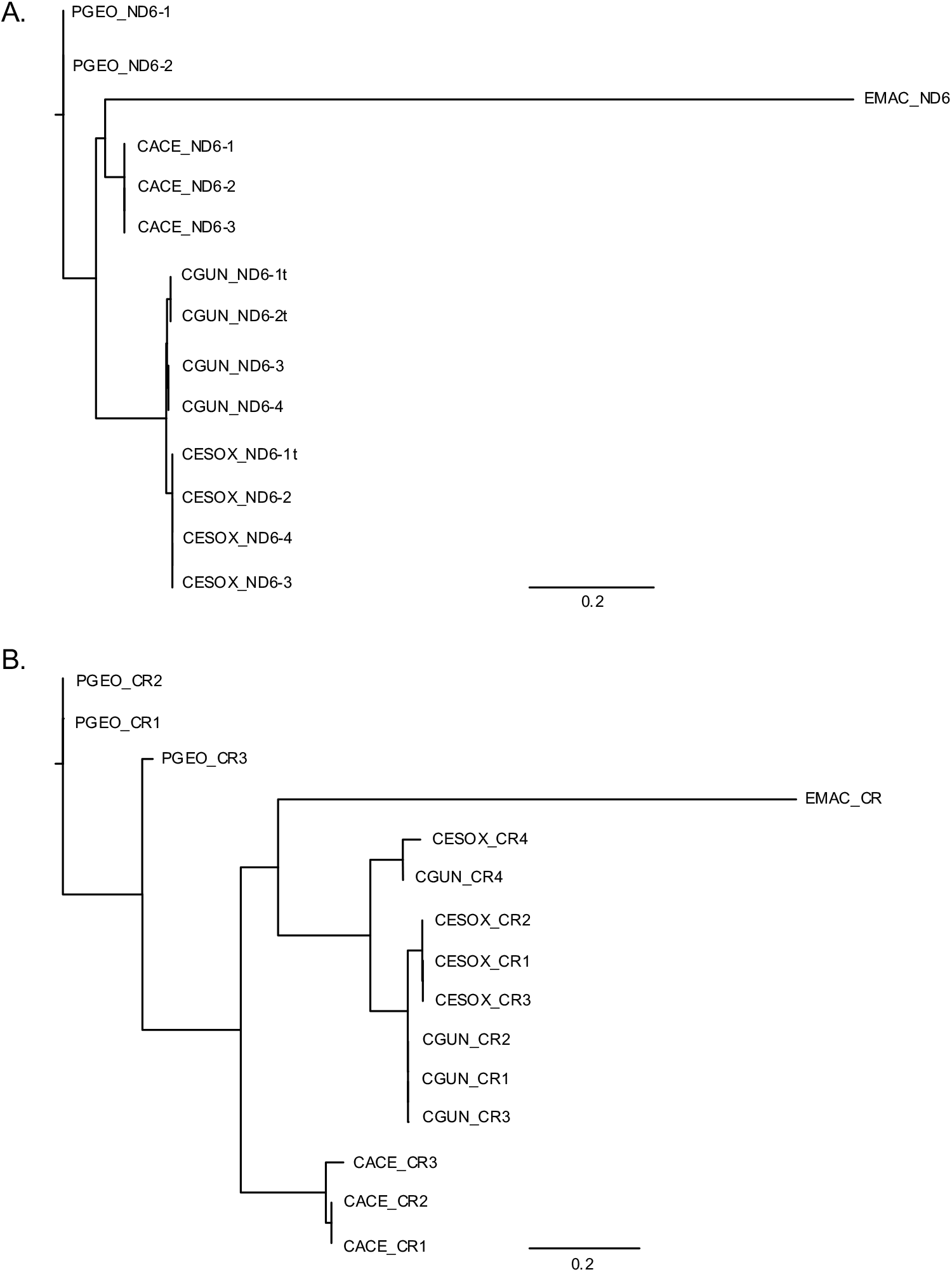
(A) Phylogenetic tree of *ND6* genes of four icefishes with *E. maclovinus* as outgroup. (B) Phylogenetic tree of control region of four icefishes with *E. maclovinus* as the outgroup.

## Discussion

### Presence of heteroplasmy could provide a reservoir for selection

Through long-read sequencing coupled with k-mer analysis, we demonstrated mt genome heteroplasmy in icefishes for the first time. The presence of more than one kind of mitogenome is often associated with aging and disease (reviewed in Elorza & Soffia, 2021; Ye et al., 2014) but our work shows that it may be more prevalent than originally thought. It is quite challenging to study the effects of heteroplasmy because it is difficult to associate phenotypic effects with different genetic copies present at low frequencies. The challenge is particularly acute since a mitogenome might not render any phenotypic effects unless it reaches a certain prevalence threshold (reviewed in Rossignol et al., 2003). The discovery of more than one mitogenomic variant in two of the four icefishes we examined demonstrates heteroplasmy, but our current approach does not tell us at what organizational level the heteroplasmy is occurring: it could occur within a specific tissue or organ, but it may also be present at a certain life stage across tissues, or the variants may be fixed within an individual but segregating across the population. A simpler reason we did not identify heteroplasmy in the remaining icefishes may simply be due to insufficient depth of mt sequencing, or the particular tissue or cell type that was used for sequencing. Mitochondrial heteroplasmy has been observed previously in humans (Payne et al., 2013), Tuatara (Macey et al., 2021), bat (Petri & Pääbo, 1996) and other species (reviewed in Barr et al., 2005) but it has been mostly limited to minor variations in length or nucleotide sequence.

For this study, we used mt reads that were sequenced incidentally with the nuclear DNA for the tissues chosen – white muscle for *C. gunnari, C. aceratus, and T. borchgrevinki*, hepatocytes (liver) for *C. esox, and spleen for P. georgianus*. Needless to say, these tissue types are quite different, with liver being the most metabolically active, followed by spleen. However, heteroplasmy was detected in *C. esox* and *C. aceratus*, demonstrating that it is occurring in disparate tissue types across at least two of the icefishes. More targeted, and tissue-specific sequencing may provide further details as to the frequency of heteroplasmy.

### Gene duplications may lead to subfunctionalization in OXPHOS Complex I

Vertebrates have evolved a very compact mitogenome, with very few intergenic sequences, and small numbers of rRNA and tRNA genes (Taanman, 1999; Wolstenholme, 1992). The conservation of small-sized vertebrate mitogenomes implies that mt gene duplications should be a rare phenomenon. However, as more mitogenomes are constructed from long reads, evidence is accumulating that mt gene duplications are prevalent (Schirtzinger et al., 2012; The Vertebrate Genomes Project Consortium et al., 2021). The formation of multiple copies of the *ND6*/*trnE*/*trnP*/CR region in four icefishes and heteroplasmy in at least two species of icefish imply that these additional copies may have functional relevance. Duplications within the nuclear genome are major contributors to adaptive evolution by generating genes which can acquire novel functions (Kondrashov, 2012). These duplicated copies are subjected to the same evolutionary forces and tend to diverge over time. In the vertebrate nuclear genome, duplicated genes are fated to pseudogenization, subfunctionalization, or neofunctionalization in rare instances, however this process is not as well understood for mt genes. Here we report possible subfunctionalization or subfunction partitioning in *ND6* in *C. esox* and *C. gunnari*.

In *C. esox* we find evidence for possible subfunction partitioning. Specifically, we identified a truncated copy of *ND6* resulting from a frameshift mutation that retains the N-terminal half of the ND6 protein. Secondary structure and TM predictions demonstrate that this is enough of the protein to maintain a wildtype N-terminal protein structure (Fig. 2F, Fig. S1). This is genetically important because the structure of the ND6 protein is highly conserved. The general structure – conserved from icefishes to zebrafish, to humans – five predicted Transmembrane domains (TMA-E). In *C. esox*, the truncation results in a wildtype structure through TMC. This in functionally important because ND6 plays several important roles in the assembly and function of the OXPHOS Complex I. ND6 functions at a very specific hinge point of Complex I helping to regulate the physical relationship between the peripheral and membrane arms. The physical relationship of these arms determines if Complex I is in a closed (active) or open (deactive) state. The transition from closed to open is partially facilitated by TMC of ND6 with the deactive state being defined by the relocation of TMD which arrests the enzyme in the deactive conformation. (Kampjut & Sazanov, 2020). Without TMD it is possible the *C. esox* truncated *ND6* copy results in a persistently closed and active state of Complex I. Interestingly, mutations almost identical to the one in *C. esox* have been functionally tested in mouse cell lines. In this case a truncation to amino acid position 79 caused by a frameshift (contrasted to position 80 in *C. esox*) resulted in decreased function of Complex I (Bai, 1998). The persistence of multiple, full copies alongside the N-terminal truncated copy of ND6 in *C. esox* could therefore serve as an evolutionary mutant model (EMM) to study Complex I activity and help understand the relationships between membrane and peripheral arm function as well as complex assembly. Furthermore, it has been hypothesized that transitions to an open (inactive) state occur at elevated temperatures (Kampjut & Sazanov, 2020). The maintenance of an ND6 protein copy incapable of switching to an open (inactive) state could therefore have been part of a *C. esox* adaptation to warmer water.

We also observed two truncations of the *ND6* gene in *C. gunnari*, each accompanied by their own set of mutations leading to alternate reading frames and different levels of total gene truncation. In one copy (T1) of *ND6*, there is a frameshift leading to only a small transcript or protein being possibly produced. It is more likely that this is evidence of pseudogenization and the RNA is degraded and never translated if it is transcribed at all. In a second copy (T2), however, a new start site and insertion of a triplet of G nucleotides results in a normal protein sequence of only the C-terminal half of ND6 (in contrast to the N-terminal half preservation in *C. esox*) (Figs. 2H, S1). Importantly, this truncation retains TME and also results in a predicted signal peptide at the N-terminal end which could be used for specifying different cellular placement of ND6 (Kapp et al., 2009). This set of mt protein truncations in *C. gunnari* seems genuinely novel, with the physiological implications unknown, providing a unique model to better understand how and if changes to Complex I, and its constituent proteins, are tied to changes in environment. Still more work is needed to confirm if the T1 and T2 truncated copies of ND6 are translated or functional.

The *ND6* gene is an important component of Complex I of the mt electron transport chain and OXPHOS pathway, and changes in its structure might lead to disease (Bai, 1998; Kishita et al., 2021). Mutations in the *ND6* gene were found to be responsible for the generation of hypoxia sensitive tumor cells in a human tumor study, and the function of ND6 within Complex I was associated with this hypoxia response (DeHaan et al., 2004). Another study (Reyes et al., 2005) has identified *ND6* as a hub of initiation of replication in chickens. The presence of duplicated copies of *ND6* in icefishes might highlight their significance in their role in surviving extreme temperatures.

### Concerted evolution

Because of the tight functional connections between protein coding genes, tRNAs, and the CR, we might expect some mt regions to evolve together by concerted evolution. Phylogenetic analysis of the duplicated *ND6* genes and CRs shows that the paralogous copies are more similar and tend to group together compared to their respective orthologs, thus suggesting concerted or parallel evolution. It also implies that duplication may have occurred independently in each species. This may imply that we fortuitously encountered these duplications before they were fully pseudogenized, which would make them an interesting case of an EMM for OXPHOS functionality. Alternatively, the gene duplications may have occurred in the common ancestor of these icefishes. It is interesting that we have seen the same block of the mitogenome duplicated to different degrees in four species of icefish – representing two separate families, which could indicate either common ancestry of this duplication, or strong selection for the functionality underlying the duplicated block.

A third possibility is that these duplications are not segregating at the population level and are therefore recent, specific, and possibly occurring frequently in the sequenced individual. Somatic copy number variations may be selectively unimportant and very common – we just have not seen them in sequencing data until now. A key question our work raises: do copy number variants reoccur in each generation, are they tissue-specific, or are they present in the germ line – eggs or sperm. Is evolution favoring distinct sets of mt for different tissues or developmental stages within an individual, or is the mt evolving at a species level in response to environmental changes, or are these variants simply the result of the drift of somatic mutations over time and therefore frequent and unremarkable. The *ND6* and CR duplication have been observed in the mt genomes of various birds, including cranes (Akiyama et al., 2017), parrots (Urantowka et al., 2018), ardeid birds (Zhou et al., 2014), and seabirds (Morris-Pocock et al., 2010). In each of these cases the *ND6*/CR duplications have been attributed to species-level, concerted evolution, which implies the presence of some form of recombination in mitogenomes, and mechanisms that might involve gene conversion events or gene turn over (Tatarenkov & Avise, 2007).

Beyond genes involved in the formation of Complex I, the presence of multiple CRs might infer a functional need to alter mt replication and transcription. Such changes could provide an advantage of increased mt genome copy number per organelle, an increased rate of replication (Jiang et al., 2007), or may play a role to increase metabolic rates in response to environmental stress (Skujina et al., 2016). Various studies have linked the presence of duplicated CRs with efficient replication mechanisms (Kumazawa et al., 1996; Umeda et al., 2001). In birds, the presence of multiple CRs is associated with longevity (Skujina et al., 2016) and confers efficient mt functionality and increased energy production required for active flight (Urantowka et al., 2018). In human cell lines that have been modified to contain mitogenomes with duplicated control regions, they were able to outcompete cell lines without the duplications (Tang et al., 2000). It has been hypothesized that multiple CRs might play a role in the survival in extreme conditions by adapting to higher energy/metabolism needs as a result of improved replication and transcription (Kinkar et al., 2020).

While the icefishes show evidence for the duplication of whole CRs, *T. borchgrevinki* contained a single, canonical CR, but its length was expanded by extensive tandem repeats. Without evolutionary pressure favoring tandem repeats, we would assume smaller mitogenomes would have an advantage because of faster replication generating more copies (Boyce et al., 1989). However, while the evolutionary significance of tandem repeats is unknown, if they are conserved in *T. borchgrevinki*, or more generally in red-blooded notothenioids, they may have a functional advantage related to better replication efficacy, improved transmission, or enhanced energy maintenance.

### Tandem repeats in *T. borchgrevinki* CR may provide a selective advantage

The increased use of long-read sequencing has enabled us to assemble the repeat repertoire in genomes with increased confidence (Kinkar et al., 2021). In the absence of any selective advantage, small mt genomes would be favored for fast replication, but the presence of larger mt genomes formed from an expanded control region (with repeats) might infer a selective advantage. Although the biological significance of these repeats is unclear, they are known to harbor elements that regulate replication/transcription. The expanded CR may be involved in better replication efficacy, improved transmission, or enhanced energy maintenance mechanisms, however more comparative work needs to be completed to understand if and to what extent these repeats are segregating in the larger population of *T. borchgrevinki* and how well these CR tandem repeats are preserved across the notothenioid clade.

### The *T. borchgrevinki* mitogenome is dominated by a large inversion

As first reported by Papetti et al., 2021, we also observed a large, inverted segment in the mt genome of *T. borchgrevinki* containing the CR along with *trnF, 12S, trnV, 16S, trnL*, *ND1* and *trnI*. Mt genomes are known to have a different base composition (either rich in G/T or alternatively A/C nucleotides) in their light and heavy strands, which results from an asymmetrical mutation process (Asakawa et al., 1991). The inversion of the CR is associated with changes in nucleotide composition of protein coding genes and is known to cause some level of reversal in compositional bias (Fonseca et al., 2014; Papetti et al., 2021). One explanation for the changes in compositional bias could be a reversal of the replication processes due to the physical inversion (and reverse complementation) of the CR. We also noticed alterations in the number of genes on the heavy and light strands compared to the canonical vertebrate mt genome. Usually, all the genes in the mt genome are on the heavy strand with only *ND6* and a few other transfer RNAs encoded on the light strand. But here the inversion resulted in the transfer of the genes in the inverted block to the strand containing *ND6*.

### Mitochondria and environmental adaptation

Mitochondria are known to have a role in the adaptation to changing environmental conditions because of their significance in important life processes (McBride et al., 2006; Sokolova, 2018). The evolution of mt genes – in response to changing energy requirements, environmental temperatures or altered oxygen proportions – can play a major role in the survival of a species since they are directly related to the formation and performance of the mt OXPHOS complexes. The power of mt genome adaptations to respond to extreme environments has been documented in cases of adaptation to high altitudes in Tibetan humans, horses, sheep and antelope, plateau pika, and Chinese snub-nosed monkeys (reviewed in Luo et al., 2013), cold temperatures (reviewed in Lubawy et al., 2022), and in other environmental stresses like temperature, hypoxia, and toxins (Sokolova, 2018) in insects. The novel genome rearrangements we observed in our study might indicate their potential role in surviving extreme temperatures. Furthermore, as described above, the presence of truncated copies of ND6 maintaining only the N-terminal half of the protein could result in persistent activation of Complex I and be an adaptation to warmer climates where Complex I is typically deactivated (Kampjut & Sazanov, 2020).

### Long-read sequencing has enabled the detection of rearrangements and structural variants

Most mitogenomes available today have been sequenced using short-read sequencing technologies which are unable to resolve complex regions containing duplications or extensive repeats (Kinkar et al., 2020; Rayamajhi et al., 2022). Our use of long-read sequencing has enabled us to assemble long tandem repeats and duplicated regions of the mitogenome and provided a platform to explore heteroplasmy. We were able to generate reads that spanned the full length of the mitochondria and in enough volume in some species to detect multiple heteroplasmic genomes in a single individual. In *C. esox* and *C. aceratus* our mt reads had an N50 length of 10,162 and 14,802 bp respectively, and the libraries in *C. aceratus* allowed us to assemble two independent genomes containing different numbers of tandemly duplicated *ND6*/*trnE*/*trnP*/CR regions. Our evidence for heteroplasmy may not reflect the full extent of the heteroplasmic conditions, as our sequencing libraries were designed to assemble nuclear genomes, and incidentally yielded only a small fraction of mt DNA reads. Long-read libraries enriched for mt DNA would be a robust approach to characterize the full magnitude of mt heteroplasmy. *T. borchgrevinki* demonstrates the utility of long-read mt libraries as the previously available short-read assemblies did not initially show the inversion, and later did not uncover the set of extensive tandem repeats in the CR (Liu et al., 2016; Papetti et al., 2021; Patel et al., 2022). Instances of mt CRs with tandem repetitive elements have been observed in various animal mt genomes (Kinkar et al., 2021), but this still might be underreported because of lack long-read-assemblies.

### K-mer analysis enabled detection of heteroplasmy

Our k-mer analysis enabled us to visualize the variable number of *ND6* copies in the tandemly duplicated *ND6*/*trnE*/*trnP*/CR region that we would have been unable to explore if we merely relied on the consensus sequence output by standard genome assemblers. For instance, for the *C. aceratus* genome, Flye generated a mitogenome assembly containing three copies of the *ND6* gene, even though a substantial number of reads (304) contained only two copies *ND6*. As Flye works by first creating a repeat graph (by collapsing repeat sequences) and then fills in the unique segments between repeat regions (Kolmogorov et al., 2019), it collapsed *ND6* and the CR. The k-mer analysis we performed could distinguish reads with two and three copies and led to the identification of heteroplasmy.

## Conclusion

Our application of long-read sequencing technology to mitochondria has highlighted a more complex genomic landscape in the mitochondria of Antarctic notothenioid fishes revealing potentially tissue- or organ-specific mitogenomes; future work must detail any functional changes resulting from the underlying heteroplasmy and determine if these genomes are reproduced somatically every generation or are part of the notothenioid germ line.

## Data Availability

The PacBio CLR raw reads for *C. gunnari, C. esox* are available from NCBI under BioProject PRJNA857989; *T. borchgrevinki* raw reads are available under BioProject PRJNA907802. The mitogenome assemblies and annotations presented in this study are hosted on Dryad.

## Acknowledgments

We thank Angel G. Rivera-Colón, and Niraj Rayamajhi for their support and feedback during our project. This work was supported by NSF OPP Grant 1645087 to JC and NSF ANT 11-42158 to C-HCC.

## References

Akiyama, T., Nishida, C., Momose, K., Onuma, M., Takami, K., & Masuda, R. (2017). Gene duplication and concerted evolution of mitochondrial DNA in crane species. Molecular Phylogenetics and Evolution, 106, 158–163. https://doi.org/10.1016/j.ympev.2016.09.026

Albertson, R. C., Cresko, W., Detrich, H. W., & Postlethwait, J. H. (2009). Evolutionary mutant models for human disease. Trends in Genetics, 25(2), 74–81. https://doi.org/10.1016/j.tig.2008.11.006

Asakawa, S., Kumazawa, Y., Araki, T., Himeno, H., Miura, K., & Watanabe, K. (1991). Strand-specific nucleotide composition bias in echinoderm and vertebrate mitochondrial genomes. Journal of Molecular Evolution, 32(6), 511–520. https://doi.org/10.1007/BF02102653

Bai, Y. (1998). The mtDNA-encoded ND6 subunit of mitochondrial NADH dehydrogenase is essential for the assembly of the membrane arm and the respiratory function of the enzyme. The EMBO Journal, 17(16), 4848–4858. https://doi.org/10.1093/emboj/17.16.4848

Barr, C. M., Neiman, M., & Taylor, D. R. (2005). Inheritance and recombination of mitochondrial genomes in plants, fungi and animals. New Phytologist, 168(1), 39–50. https://doi.org/10.1111/j.1469-8137.2005.01492.x

Beck, E. A., Healey, H. M., Small, C. M., Currey, M. C., Desvignes, T., Cresko, W. A., & Postlethwait, J. H. (2022). Advancing human disease research with fish evolutionary mutant models. Trends in Genetics, 38(1), 22–44. https://doi.org/10.1016/j.tig.2021.07.002

Benson, G. (1999). Tandem repeats finder: A program to analyze DNA sequences. Nucleic Acids Research, 27(2), 573–580. https://doi.org/10.1093/nar/27.2.573

Bilyk, K. T., & DeVries, A. L. (2011). Heat tolerance and its plasticity in Antarctic fishes. Comparative Biochemistry and Physiology Part A: Molecular & Integrative Physiology, 158(4), 382–390. https://doi.org/10.1016/j.cbpa.2010.12.010

Bista, I., Wood, J. M. D., Desvignes, T., McCarthy, S. A., Matschiner, M., Ning, Z., Tracey, A., Torrance, J., Sims, Y., Chow, W., Smith, M., Oliver, K., Haggerty, L., Salzburger, W., Postlethwait, J. H., Howe, K., Clark, M. S., Detrich, W. H., Cheng, C.-H. C., … Durbin, R. (2022). Genomics of cold adaptations in the Antarctic notothenioid fish radiation. BioRxiv, 2022.06.08.494096. https://doi.org/10.1101/2022.06.08.494096

Boore, J. L. (1999). Animal mitochondrial genomes. Nucleic Acids Research, 27(8), 1767–1780. https://doi.org/10.1093/nar/27.8.1767

Boyce, T. M., Zwick, M. E., & Aquadro, C. F. (1989). Mitochondrial DNA in the bark weevils: Size, structure and heteroplasmy. Genetics, 123(4), 825–836. https://doi.org/10.1093/genetics/123.4.825

Breton, S., & Stewart, D. T. (2015). Atypical mitochondrial inheritance patterns in eukaryotes. Genome, 58(10), 423–431. https://doi.org/10.1139/gen-2015-0090

Brierley, E. J., Johnson, M. A., Lightowlers, R. N., James, O. F. W., & Turnbull, D. M. (1998). Role of mitochondrial DNA mutations in human aging: Implications for the central nervous system and muscle. Annals of Neurology, 43(2), 217–223. https://doi.org/10.1002/ana.410430212

Broughton, R. E., Milam, J. E., & Roe, B. A. (2001). The Complete Sequence of the Zebrafish (*Danio rerio*) Mitochondrial Genome and Evolutionary Patterns in Vertebrate Mitochondrial DNA. Genome Research, 11(11), 1958–1967. https://doi.org/10.1101/gr.156801

Camacho, C., Coulouris, G., Avagyan, V., Ma, N., Papadopoulos, J., Bealer, K., & Madden, T. L. (2009). BLAST+: Architecture and applications. BMC Bioinformatics, 10(1), 421. https://doi.org/10.1186/1471-2105-10-421

Chan, P. P., Lin, B. Y., Mak, A. J., & Lowe, T. M. (2021). tRNAscan-SE 2.0: Improved detection and functional classification of transfer RNA genes. Nucleic Acids Research, 49(16), 9077–9096. https://doi.org/10.1093/nar/gkab688

Chinnery, P. F., Brown, D. T., Andrews, R. M., Singh-Kler, R., Riordan-Eva, P., Lindley, J., Applegarth, D. A., Turnbull, D. M., & Howell, N. (2001). The mitochondrial ND6 gene is a hot spot for mutations that cause Leber’s hereditary optic neuropathy. Brain, 124(1), 209–218. https://doi.org/10.1093/brain/124.1.209

Cocca, E., Ratnayake-Lecamwasam, M., Parker, S. K., Camardella, L., Ciaramella, M., di Prisco, G., & Detrich, H. W. (1995). Genomic remnants of alpha-globin genes in the hemoglobinless antarctic icefishes. Proceedings of the National Academy of Sciences, 92(6), 1817–1821. https://doi.org/10.1073/pnas.92.6.1817

Cunningham, F., Allen, J. E., Allen, J., Alvarez-Jarreta, J., Amode, M. R., Armean, I. M., Austine-Orimoloye, O., Azov, A. G., Barnes, I., Bennett, R., Berry, A., Bhai, J., Bignell, A., Billis, K., Boddu, S., Brooks, L., Charkhchi, M., Cummins, C., Da Rin Fioretto, L., … Flicek, P. (2022). Ensembl 2022. Nucleic Acids Research, 50(D1), D988–D995. https://doi.org/10.1093/nar/gkab1049

DeHaan, C., Habibi-Nazhad, B., Yan, E., Salloum, N., Parliament, M., & Allalunis-Turner, J. (2004). Mutation in mitochondrial complex I ND6 subunit is associated with defective response to hypoxia in human glioma cells. Molecular Cancer, 3(1), 19. https://doi.org/10.1186/1476-4598-3-19

DeVries, A. L., & Cheng, C. -H. C. (2005). Antifreeze Proteins and Organismal Freezing Avoidance in Polar Fishes. In Fish Physiology (Vol. 22, pp. 155–201). Elsevier. https://doi.org/10.1016/S1546-5098(04)22004-0

Donath, A., Jühling, F., Al-Arab, M., Bernhart, S. H., Reinhardt, F., Stadler, P. F., Middendorf, M., & Bernt, M. (2019). Improved annotation of protein-coding genes boundaries in metazoan mitochondrial genomes. Nucleic Acids Research, 47(20), 10543–10552. https://doi.org/10.1093/nar/gkz833

Eastman, J. T. (1993). Antarctic fish biology: Evolution in a unique environment. Academic Press.

Eastman, J. T. (2005). The nature of the diversity of Antarctic fishes. Polar Biology, 28(2), 93–107. https://doi.org/10.1007/s00300-004-0667-4

Edgar, R. C. (2004). MUSCLE: Multiple sequence alignment with high accuracy and high throughput. Nucleic Acids Research, 32(5), 1792–1797. https://doi.org/10.1093/nar/gkh340

Elorza, A. A., & Soffia, J. P. (2021). mtDNA Heteroplasmy at the Core of Aging-Associated Heart Failure. An Integrative View of OXPHOS and Mitochondrial Life Cycle in Cardiac Mitochondrial Physiology. Frontiers in Cell and Developmental Biology, 9, 625020. https://doi.org/10.3389/fcell.2021.625020

Fitch, N. A., Johnson, I. A., & Wood, R. E. (1984). Skeletal muscle capillary supply in a fish that lacks respiratory pigments. Respiration Physiology, 57(2), 201–211. https://doi.org/10.1016/0034-5687(84)90093-8

Fonseca, M. M., Harris, D. J., & Posada, D. (2014). The Inversion of the Control Region in Three Mitogenomes Provides Further Evidence for an Asymmetric Model of Vertebrate mtDNA Replication. PLoS ONE, 9(9), e106654. https://doi.org/10.1371/journal.pone.0106654

Fontanesi, F. (2015). Mitochondria: Structure and Role in Respiration. In John Wiley & Sons, Ltd (Ed.), ELS (1st ed., pp. 1–13). Wiley. https://doi.org/10.1002/9780470015902.a0001380.pub2

Force, A., Lynch, M., Pickett, F. B., Amores, A., Yan, Y., & Postlethwait, J. (1999). Preservation of Duplicate Genes by Complementary, Degenerative Mutations. Genetics, 151(4), 1531–1545. https://doi.org/10.1093/genetics/151.4.1531

Gong, L., Shi, W., Wang, Z.-M., Miao, X.-G., & Kong, X.-Y. (2013). Control region translocation and a tRNA gene inversion in the mitogenome of *Paraplagusia japonica* (Pleuronectiformes: Cynoglossidae). Mitochondrial DNA, 24(6), 671–673. https://doi.org/10.3109/19401736.2013.773984

Guindon, S., Dufayard, J.-F., Lefort, V., Anisimova, M., Hordijk, W., & Gascuel, O. (2010). New Algorithms and Methods to Estimate Maximum-Likelihood Phylogenies: Assessing the Performance of PhyML 3.0. Systematic Biology, 59(3), 307–321. https://doi.org/10.1093/sysbio/syq010

Haag-Liautard, C., Coffey, N., Houle, D., Lynch, M., Charlesworth, B., & Keightley, P. D. (2008). Direct Estimation of the Mitochondrial DNA Mutation Rate in Drosophila melanogaster. PLoS Biology, 6(8), e204. https://doi.org/10.1371/journal.pbio.0060204

Hallgren, J., Tsirigos, K. D., Pedersen, M. D., Almagro Armenteros, J. J., Marcatili, P., Nielsen, H., Krogh, A., & Winther, O. (2022). DeepTMHMM predicts alpha and beta transmembrane proteins using deep neural networks. BioRxiv, 2022.04.08.487609. https://doi.org/10.1101/2022.04.08.487609

Hemmingsen, E. A., Douglas, E. L., Johansen, K., & Millard, R. W. (1972). Aortic blood flow and cardiac output in the hemoglobin-free fish Chaenocephalus aceratus. Comparative Biochemistry and Physiology Part A: Physiology, 43(4), 1045–1051. https://doi.org/10.1016/0300-9629(72)90176-4

Hommelsheim, C. M., Frantzeskakis, L., Huang, M., & Ülker, B. (2015). PCR amplification of repetitive DNA: A limitation to genome editing technologies and many other applications. Scientific Reports, 4(1), 5052. https://doi.org/10.1038/srep05052

Inoue, J. G. (2003). Evolution of the Deep-Sea Gulper Eel Mitochondrial Genomes: Large-Scale Gene Rearrangements Originated Within the Eels. Molecular Biology and Evolution, 20(11), 1917–1924. https://doi.org/10.1093/molbev/msg206

Irwin, J. A., Saunier, J. L., Niederstätter, H., Strouss, K. M., Sturk, K. A., Diegoli, T. M., Brandstätter, A., Parson, W., & Parsons, T. J. (2009). Investigation of Heteroplasmy in the Human Mitochondrial DNA Control Region: A Synthesis of Observations from More Than 5000 Global Population Samples. Journal of Molecular Evolution, 68(5), 516–527. https://doi.org/10.1007/s00239-009-9227-4

Itsara, L. S., Kennedy, S. R., Fox, E. J., Yu, S., Hewitt, J. J., Sanchez-Contreras, M., Cardozo-Pelaez, F., & Pallanck, L. J. (2014). Oxidative Stress Is Not a Major Contributor to Somatic Mitochondrial DNA Mutations. PLoS Genetics, 10(2), e1003974. https://doi.org/10.1371/journal.pgen.1003974

Jiang, Z. J., Castoe, T. A., Austin, C. C., Burbrink, F. T., Herron, M. D., McGuire, J. A., Parkinson, C. L., & Pollock, D. D. (2007). Comparative mitochondrial genomics of snakes: Extraordinary substitution rate dynamics and functionality of the duplicate control region. BMC Evolutionary Biology, 7(1), 123. https://doi.org/10.1186/1471-2148-7-123

Jin, W.-T., Guan, J.-Y., Dai, X.-Y., Wu, G.-J., Zhang, L.-P., Storey, K. B., Zhang, J.-Y., Zheng, R.-Q., & Yu, D.-N. (2022). Mitochondrial gene expression in different organs of Hoplobatrachus rugulosus from China and Thailand under low-temperature stress. BMC Zoology, 7(1), 24. https://doi.org/10.1186/s40850-022-00128-7

Johnston, I. A., Calvo, J., Guderley, H., Fernandez, D., & Palmer, L. (1998). Latitudinal variation in the abundance and oxidative capacities of muscle mitochondria in perciform fishes. Journal of Experimental Biology, 201(1), 1–12. https://doi.org/10.1242/jeb.201.1.1

Kampjut, D., & Sazanov, L. A. (2020). The coupling mechanism of mammalian respiratory complex I. Science, 370 (6516), eabc4209. https://doi.org/10.1126/science.abc4209

Kapp, K., Schrempf, S., Lemberg, M. K., & Dobberstein, B. (2009). Post-Targeting Functions of Signal Peptides. In Protein transport into the endoplasmic reticulum (pp. 1–16).

Kim, B.-M., Amores, A., Kang, S., Ahn, D.-H., Kim, J.-H., Kim, I.-C., Lee, J. H., Lee, S. G., Lee, H., Lee, J., Kim, H.-W., Desvignes, T., Batzel, P., Sydes, J., Titus, T., Wilson, C. A., Catchen, J. M., Warren, W. C., Schartl, M., … Park, H. (2019). Antarctic blackfin icefish genome reveals adaptations to extreme environments. Nature Ecology & Evolution, 3(3), 469–478. https://doi.org/10.1038/s41559-019-0812-7

Kinkar, L., Gasser, R., Webster, B., Rollinson, D., Littlewood, D., Chang, B., Stroehlein, A., Korhonen, P., & Young, N. (2021). Nanopore Sequencing Resolves Elusive Long Tandem-Repeat Regions in Mitochondrial Genomes. International Journal of Molecular Sciences, 22(4), 1811. https://doi.org/10.3390/ijms22041811

Kinkar, L., Young, N. D., Sohn, W.-M., Stroehlein, A. J., Korhonen, P. K., & Gasser, R. B. (2020). First record of a tandem-repeat region within the mitochondrial genome of Clonorchis sinensis using a long-read sequencing approach. PLOS Neglected Tropical Diseases, 14(8), e0008552. https://doi.org/10.1371/journal.pntd.0008552

Kishita, Y., Ishikawa, K., Nakada, K., Hayashi, J.-I., Fushimi, T., Shimura, M., Kohda, M., Ohtake, A., Murayama, K., & Okazaki, Y. (2021). A high mutation load of m.14597A>G in MT-ND6 causes Leigh syndrome. Scientific Reports, 11(1), 11123. https://doi.org/10.1038/s41598-021-90196-5

Klucnika, A., & Ma, H. (2019). A battle for transmission: The cooperative and selfish animal mitochondrial genomes. Open Biology, 9(3), 180267. https://doi.org/10.1098/rsob.180267

Kock, K.-H. (2005). Antarctic icefishes (Channichthyidae): A unique family of fishes. A review, Part I. Polar Biology, 28(11), 862–895. https://doi.org/10.1007/s00300-005-0019-z

Kolmogorov, M., Yuan, J., Lin, Y., & Pevzner, P. A. (2019). Assembly of long, error-prone reads using repeat graphs. Nature Biotechnology, 37(5), 540–546. https://doi.org/10.1038/s41587-019-0072-8

Kondrashov, F. A. (2012). Gene duplication as a mechanism of genomic adaptation to a changing environment. Proceedings of the Royal Society B: Biological Sciences, 279(1749), 5048–5057. https://doi.org/10.1098/rspb.2012.1108

Kong, X., Dong, X., Zhang, Y., Shi, W., Wang, Z., & Yu, Z. (2009). A novel rearrangement in the mitochondrial genome of tongue sole, *Cynoglossus semilaevis*: Control region translocation and a tRNA gene inversion. Genome, 52(12), 975–984. https://doi.org/10.1139/G09-069

Koren, S., Walenz, B. P., Berlin, K., Miller, J. R., Bergman, N. H., & Phillippy, A. M. (2017). Canu: Scalable and accurate long-read assembly via adaptive *k* -mer weighting and repeat separation. Genome Research, 27(5), 722–736. https://doi.org/10.1101/gr.215087.116

Kumazawa, Y., Ota, H., Nishida, M., & Ozawa, T. (1996). Gene rearrangements in snake mitochondrial genomes: Highly concerted evolution of control-region-like sequences duplicated and inserted into a tRNA gene cluster. Molecular Biology and Evolution, 13(9), 1242–1254. https://doi.org/10.1093/oxfordjournals.molbev.a025690

Lee, W.-J., Conroy, J., Howell, W. H., & Kocher, ThomasD. (1995). Structure and evolution of teleost mitochondrial control regions. Journal of Molecular Evolution, 41(1), 54–66. https://doi.org/10.1007/BF00174041

Leister, D. (2005). Origin, evolution and genetic effects of nuclear insertions of organelle DNA. Trends in Genetics, 21(12), 655–663. https://doi.org/10.1016/j.tig.2005.09.004

Li, H. (2018). Minimap2: Pairwise alignment for nucleotide sequences. Bioinformatics, 34(18), 3094–3100. https://doi.org/10.1093/bioinformatics/bty191

Liu, Y., Yang, M., Zhou, T., Xing, H., Chen, L., & Zhang, D. (2016). Complete mitochondrial genome of the Antarctic cod icefish, *Pagothenia borchgrevinki* (Perciformes: Nototheniidae). Mitochondrial DNA Part B, 1(1), 432–433. https://doi.org/10.1080/23802359.2016.1180554

Lubawy, J., Chowański, S., Adamski, Z., & Słocińska, M. (2022). Mitochondria as a target and central hub of energy division during cold stress in insects. Frontiers in Zoology, 19(1), 1. https://doi.org/10.1186/s12983-021-00448-3

Luo, Y., Yang, X., & Gao, Y. (2013). Mitochondrial DNA response to high altitude: A new perspective on high-altitude adaptation. Mitochondrial DNA, 24(4), 313–319. https://doi.org/10.3109/19401736.2012.760558

Macey, J. R., Pabinger, S., Barbieri, C. G., Buring, E. S., Gonzalez, V. L., Mulcahy, D. G., DeMeo, D. P., Urban, L., Hime, P. M., Prost, S., Elliott, A. N., & Gemmell, N. J. (2021). Evidence of two deeply divergent co-existing mitochondrial genomes in the Tuatara reveals an extremely complex genomic organization. Communications Biology, 4(1), 116. https://doi.org/10.1038/s42003-020-01639-0

McBride, H. M., Neuspiel, M., & Wasiak, S. (2006). Mitochondria: More Than Just a Powerhouse. Current Biology, 16(14), R551–R560. https://doi.org/10.1016/j.cub.2006.06.054

Mignotte, F., Gueride, M., Champagne, A.-M., & Mounolou, J.-C. (1990). Direct repeats in the non-coding region of rabbit mitochondrial DNA. Involvement in the generation of intra-and inter-individual heterogeneity. European Journal of Biochemistry, 194(2), 561–571. https://doi.org/10.1111/j.1432-1033.1990.tb15653.x

Miya, M., & Nishida, M. (1999). Organization of the Mitochondrial Genome of a Deep-Sea Fish, Gonostoma gracile (Teleostei: Stomiiformes): First Example of Transfer RNA Gene Rearrangements in Bony Fishes. Marine Biotechnology, 1(5), 416–426. https://doi.org/10.1007/PL00011798

Morris-Pocock, J. A., Taylor, S. A., Birt, T. P., & Friesen, V. L. (2010). Concerted evolution of duplicated mitochondrial control regions in three related seabird species. BMC Evolutionary Biology, 10(1), 14. https://doi.org/10.1186/1471-2148-10-14

Near, T. J., MacGuigan, D. J., Parker, E., Struthers, C. D., Jones, C. D., & Dornburg, A. (2018). Phylogenetic analysis of Antarctic notothenioids illuminates the utility of RADseq for resolving Cenozoic adaptive radiations. Molecular Phylogenetics and Evolution, 129, 268–279. https://doi.org/10.1016/j.ympev.2018.09.001

O’Brien, K. M., & Mueller, I. A. (2010). The Unique Mitochondrial Form and Function of Antarctic Channichthyid Icefishes. Integrative and Comparative Biology, 50(6), 993–1008. https://doi.org/10.1093/icb/icq038

O’Brien, K. M., & Sidell, B. D. (2000). The interplay among cardiac ultrastructure, metabolism and the expression of oxygen-binding proteins in Antarctic fishes. Journal of Experimental Biology, 203(8), 1287–1297. https://doi.org/10.1242/jeb.203.8.1287

Ojala, D., Montoya, J., & Attardi, G. (1981). TRNA punctuation model of RNA processing in human mitochondria. Nature, 290(5806), 470–474. https://doi.org/10.1038/290470a0

Papetti, C., Babbucci, M., Dettai, A., Basso, A., Lucassen, M., Harms, L., Bonillo, C., Heindler, F. M., Patarnello, T., & Negrisolo, E. (2021). Not Frozen in the Ice: Large and Dynamic Rearrangements in the Mitochondrial Genomes of the Antarctic Fish. Genome Biology and Evolution, 13(3), evab017. https://doi.org/10.1093/gbe/evab017

Parakatselaki, M.-E., & Ladoukakis, E. D. (2021). mtDNA Heteroplasmy: Origin, Detection, Significance, and Evolutionary Consequences. Life, 11(7), 633. https://doi.org/10.3390/life11070633

Patel, S., Evans, C. W., Stuckey, A., Matzke, N. J., & Millar, C. D. (2022). A Unique Mitochondrial Gene Block Inversion in Antarctic Trematomin Fishes: A Cautionary Tale. Journal of Heredity, 113(4), 414–420. https://doi.org/10.1093/jhered/esac028

Payne, B. A. I., Wilson, I. J., Yu-Wai-Man, P., Coxhead, J., Deehan, D., Horvath, R., Taylor, R. W., Samuels, D. C., Santibanez-Koref, M., & Chinnery, P. F. (2013). Universal heteroplasmy of human mitochondrial DNA. Human Molecular Genetics, 22(2), 384–390. https://doi.org/10.1093/hmg/dds435

Pereira, S. L. (2000). Mitochondrial genome organization and vertebrate phylogenetics. Genetics and Molecular Biology, 23(4), 745–752. https://doi.org/10.1590/S1415-47572000000400008

Petri, B., & Pääbo, S. (1996). Extreme Sequence Heteroplasmy in Bat Mitochondria! DNA. 8.

Ramos, E. K. da S., Freitas, L., & Nery, M. F. (2020). The role of selection in the evolution of marine turtles mitogenomes. Scientific Reports, 10(1), 16953. https://doi.org/10.1038/s41598-020-73874-8

Rayamajhi, N., Cheng, C.-H. C., & Catchen, J. M. (2022). Evaluating Illumina-, Nanopore-, and PacBio-based genome assembly strategies with the bald notothen, *Trematomus borchgrevinki*. G3 Genes|Genomes|Genetics, jkac192. https://doi.org/10.1093/g3journal/jkac192

Reuter, J. S., & Mathews, D. H. (2010). RNAstructure: Software for RNA secondary structure prediction and analysis. BMC Bioinformatics, 11(1), 129. https://doi.org/10.1186/1471-2105-11-129

Reyes, A., Yang, M. Y., Bowmaker, M., & Holt, I. J. (2005). Bidirectional Replication Initiates at Sites Throughout the Mitochondrial Genome of Birds. Journal of Biological Chemistry, 280(5), 3242–3250. https://doi.org/10.1074/jbc.M411916200

Rivera-Colón, A. G., Rayamajhi, N., Minhas, B. F., Madrigal, G., Bilyk, K. T., Yoon, V., Hüne, M., Gregory, S., Cheng, C.-H. C., & Catchen, J. M. (2022). Genomics of Secondarily Temperate Adaptation in the Only Non-Antarctic Icefish. BioRxiv, 2022.08.13.503862. https://doi.org/10.1101/2022.08.13.503862

Rossignol, R., Faustin, B., Rocher, C., Malgat, M., Mazat, J.-P., & Letellier, T. (2003). Mitochondrial threshold effects. Biochemical Journal, 370(3), 751–762. https://doi.org/10.1042/bj20021594

Ruud, J. T. (1954). Vertebrates without Erythrocytes and Blood Pigment. Nature, 173(4410), 848–850. https://doi.org/10.1038/173848a0

Sagan, L. (1967). On the origin of mitosing cells. Journal of Theoretical Biology, 14(3), 225–IN6. https://doi.org/10.1016/0022-5193(67)90079-3

Satoh, T. P., Miya, M., Mabuchi, K., & Nishida, M. (2016). Structure and variation of the mitochondrial genome of fishes. BMC Genomics, 17(1), 719. https://doi.org/10.1186/s12864-016-3054-y

Schirtzinger, E. E., Tavares, E. S., Gonzales, L. A., Eberhard, J. R., Miyaki, C. Y., Sanchez, J. J., Hernandez, A., Müeller, H., Graves, G. R., Fleischer, R. C., & Wright, T. F. (2012). Multiple independent origins of mitochondrial control region duplications in the order Psittaciformes. Molecular Phylogenetics and Evolution, 64(2), 342–356. https://doi.org/10.1016/j.ympev.2012.04.009

Shi, W., Dong, X.-L., Wang, Z.-M., Miao, X.-G., Wang, S.-Y., & Kong, X.-Y. (2013). Complete mitogenome sequences of four flatfishes (Pleuronectiformes) reveal a novel gene arrangement of L-strand coding genes. BMC Evolutionary Biology, 13(1), 173. https://doi.org/10.1186/1471-2148-13-173

Sidell, B. D., & O’Brien, K. M. (2006). When bad things happen to good fish: The loss of hemoglobin and myoglobin expression in Antarctic icefishes. Journal of Experimental Biology, 209(10), 1791–1802. https://doi.org/10.1242/jeb.02091

Skujina, I., McMahon, R., Lenis, V. P. E., Gkoutos, G. V., & Hegarty, M. (2016). Duplication of the mitochondrial control region is associated with increased longevity in birds. Aging, 8(8), 1781–1789. https://doi.org/10.18632/aging.101012

Sokolova, I. (2018). Mitochondrial Adaptations to Variable Environments and Their Role in Animals’ Stress Tolerance. Integrative and Comparative Biology, 58(3), 519–531. https://doi.org/10.1093/icb/icy017

Stankovic, A., Spalik, K., Kamler, E., Borsuk, P., & Weglenski, P. (2002). Recent origin of sub-Antarctic notothenioids. Polar Biology, 25(3), 203–205. https://doi.org/10.1007/s00300-001-0327-x

Stewart, J. B., & Chinnery, P. F. (2015). The dynamics of mitochondrial DNA heteroplasmy: Implications for human health and disease. Nature Reviews Genetics, 16(9), 530–542. https://doi.org/10.1038/nrg3966

Taanman, J.-W. (1999). The mitochondrial genome: Structure, transcription, translation and replication. Biochimica et Biophysica Acta (BBA) - Bioenergetics, 1410(2), 103–123. https://doi.org/10.1016/S0005-2728(98)00161-3

Tang, Y., Manfredi, G., Hirano, M., & Schon, E. A. (2000). Maintenance of Human Rearranged Mitochondrial DNAs in Long-Term Cultured Transmitochondrial Cell Lines. Molecular Biology of the Cell, 11(7), 2349–2358. https://doi.org/10.1091/mbc.11.7.2349

Tatarenkov, A., & Avise, J. C. (2007). Rapid concerted evolution in animal mitochondrial DNA. Proceedings of the Royal Society B: Biological Sciences, 274(1619), 1795–1798. https://doi.org/10.1098/rspb.2007.0169

The Vertebrate Genomes Project Consortium, Formenti, G., Rhie, A., Balacco, J., Haase, B., Mountcastle, J., Fedrigo, O., Brown, S., Capodiferro, M. R., Al-Ajli, F. O., Ambrosini, R., Houde, P., Koren, S., Oliver, K., Smith, M., Skelton, J., Betteridge, E., Dolucan, J., Corton, C., … Jarvis, E. D. (2021). Complete vertebrate mitogenomes reveal widespread repeats and gene duplications. Genome Biology, 22(1), 120. https://doi.org/10.1186/s13059-021-02336-9

Umeda, S., Tang, Y., Okamoto, M., Hamasaki, N., Schon, E. A., & Kang, D. (2001). Both Heavy Strand Replication Origins Are Active in Partially Duplicated Human Mitochondrial DNAs. Biochemical and Biophysical Research Communications, 286(4), 681–687. https://doi.org/10.1006/bbrc.2001.5436

Urantowka, A. D., Kroczak, A., Silva, T., Padrón, R. Z., Gallardo, N. F., Blanch, J., Blanch, B., & Mackiewicz, P. (2018). New insight into parrots’ mitogenomes indicates that their ancestor contained a duplicated region. Molecular Biology and Evolution, 35(12), 2989–3009. https://doi.org/10.1093/molbev/msy189

Wolstenholme, D. R. (1992). Animal Mitochondrial DNA: Structure and Evolution. In International Review of Cytology (Vol. 141, pp. 173–216). Elsevier. https://doi.org/10.1016/S0074-7696(08)62066-5

Xu, S., Luosang, J., Hua, S., He, J., Ciren, A., Wang, W., Tong, X., Liang, Y., Wang, J., & Zheng, X. (2007). High Altitude Adaptation and Phylogenetic Analysis of Tibetan Horse Based on the Mitochondrial Genome. Journal of Genetics and Genomics, 34(8), 720–729. https://doi.org/10.1016/S1673-8527(07)60081-2

Ye, K., Lu, J., Ma, F., Keinan, A., & Gu, Z. (2014). Extensive pathogenicity of mitochondrial heteroplasmy in healthy human individuals. Proceedings of the National Academy of Sciences, 111(29), 10654–10659. https://doi.org/10.1073/pnas.1403521111

Zhou, X., Lin, Q., Fang, W., & Chen, X. (2014). The complete mitochondrial genomes of sixteen ardeid birds revealing the evolutionary process of the gene rearrangements. BMC Genomics, 15(1), 573. https://doi.org/10.1186/1471-2164-15-573

