## Supplementary Figure 1 for "Novel mitochondrial genome rearrangements including duplications and extensive heteroplasmy could underlie temperature adaptations in Antarctic Notothenioid Fishes"

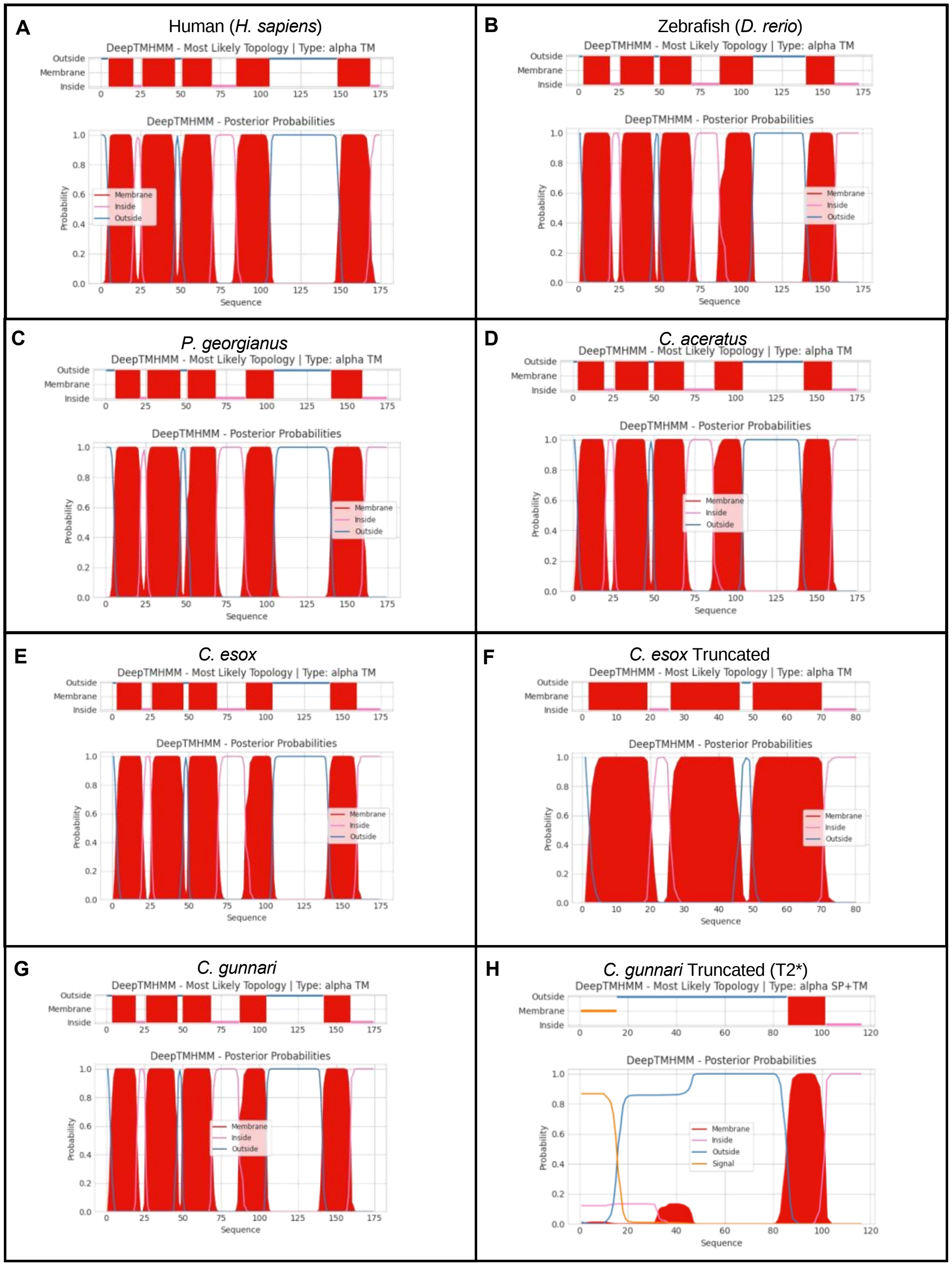


**Figure S1.** DeepTMHMM transmembrane predictions for ND6 protein in (A) Human, (B) Zebrafish, (C) *P. georgianus*, (D) *C. aceratus,* (E) *C.esox,* (F) *C.esox truncated,* (G) *C. gunnari,* (H) *C.gunnari truncated (T2)*
